# RCoxNet: deep learning framework for enhanced cancer survival prediction integrating random walk with restart with mutation and clinical data

**DOI:** 10.1101/2024.09.17.613428

**Authors:** Stuti Kumari, Sakshi Gujral, Smruti Panda, Prashant Gupta, Gaurav Ahuja, Debarka Sengupta

## Abstract

Cancer poses a significant global health challenge, characterized by a complex disease progression and disrupted growth regulation. A thorough understanding of cellular and molecular biological mechanisms is essential for developing novel treatments and improving the accuracy of patient survival predictions. While prior studies have leveraged gene expression and clinical data to forecast survival outcomes through current machine learning and deep learning approaches, gene mutation data—despite being a widely recognized metric—has rarely been incorporated due to its limited information, inadequate representation of gene relationships, and data sparsity, which negatively affects the robustness, effectiveness, and interpretability of current survival analysis approaches. To overcome the challenges of mutation data sparsity, we propose RCoxNet, a novel deep learning neural network framework that integrates the Random Walk with Restart (RWR) algorithm with a deep learning Cox Proportional Hazards model. By applying this framework to mutation data from cBioportal, our model achieved an average concordance index of 0.62 ± 0.05 across four cancer types, outperforming existing deep neural network models. Additionally, we identified clinical features critical for differentiating between predicted high- and low-risk patients, with the relevance of these features being partially supported by previous studies.

## 1. Introduction

Cancer remains a leading global cause of mortality, prompting extensive medical research dedicated to the survival analysis of afflicted patients. This is initiated by genetic and epigenetic changes within specific cells, some of which have the potential to spread and infiltrate other tissues. While the processes involved in carcinogenesis and the progression of tumors are diverse, three key aspects are proved to be crucial in tumor biology: genomic and epigenomic modifications driving cell transformation, the specific cell types undergoing these alterations, and the mechanisms of invasion and metastasis, which significantly influence the aggressiveness of tumors (Sánchez P et al. 2021). In the last decade, advancements in genomics and epigenomics technologies have yielded vast amounts of data aimed at unraveling the intricate mechanisms underlying cancer. Predicted survival risk plays a pivotal role in guiding treatment decisions influencing the choice and assessment of treatment methods (Roy G et al. 2022). Accurate cancer prognosis prediction helps clinicians to conduct more appropriate treatment allocation for patients to prolong life span, increase life quality, and reduce unnecessary treatment cost. Recent studies have applied machine learning (ML) techniques in the analysis of clinical and genomic features, and they showed that ML has improved performance in cancer susceptibility, recurrence, and survival prediction compared to traditional approaches (e.g., Kaplan–Meier method). In practice, several issues can undermine the robustness of survival predictions. Firstly, measuring some important clinical variables relies heavily on the clinician’s individual interpretation, which may introduce human bias, thereby reducing the accuracy and reliability of the prediction results. Secondly, small sample size accompanied by high-dimensional input data (e.g., gene expression data, whole slide image, etc.) can result in overfitting and hamper the generalizability of existing models. Thirdly, the relationship between predictors and survival outcome may be non-linear, and thus existing models that assume a linear relationship (e.g., Cox Proportional Hazards model) may produce inaccurate results. As a trending approach, DNN can also deal with non-linear relationships intrinsically, which can well represent complicated data structures. Owing to their flexibility, deep learning (DL) models can be designed (or combined with other approaches) to conduct feature extraction and integration from high-dimensional omics data. Deepsurv is a model that is based on CoxPH but adopts a DNN structure. Katzman et al. demonstrated that Deepsurv was able to outperform the CoxPH model in prognosis prediction under various scenarios, highlighting the strength of deep learning models in handling complex data patterns compared to conventional approaches. In a similar fashion, Cox-nnet adopted feed-forward DNN structures for prognosis prediction, but unlike DeepSURV, Coxnnet (Ching T. et al., 2018), AECOX (Huang Z. et al., 2020), and SurvivalNet (Yousefi S. et al., 2017) can handle high-dimensional gene expression data. Current approaches incorporate knowledge of gene pathways alongside mutation data, but they face reliability issues, particularly concerning the sparsity of data. These methods encounter challenges in effectively handling sparse data due to the inclusion of gene pathways. Additionally, existing methodologies for survival analysis may suffer from limited robustness, interpretability, and the ability to capture complex relationships within the data, potentially hindering their efficacy in practical applications (Liu J et al. 2023, Sun B et al. 2023).

In recent years, there has been a notable increase in the accumulation of physical and functional interactions among biological macromolecules. One of the widely used concepts to understand the network structure of PPI networks is random walks formulated to investigate the overall structure of networks. This involves simulating a particle that systematically traverses from one node to a randomly chosen neighboring node (Valdeolivas A et al. 2019, Huan T et al. 2014). While existing research has not demonstrated much about the significant impact of tumor mutation burden (TMB) and microsatellite instability (MSI) with interaction networks, it remains challenging to involve these clinical variables. Tumor Mutation Burden (TMB) refers to the total number of mutations identified in a tumor’s genome, which serves as a biomarker to predict the effectiveness of therapy for specific cancer types. TMB is thought to play a crucial role in generating immunogenic neo-peptides presented on major histocompatibility complexes (MHC) on the surface of tumor cells, influencing patient responses to Immune Checkpoint Inhibitors (Sha D et al. 2020). Microsatellite Instability (MSI) is another predictive cancer biomarker that occurs when a short repeated sequence of DNA (microsatellite or short tandem repeat) undergoes a mutation during replication (Strickler JH et al. 2021, Chakrabarti S et al. 2022). Precise detection of MSI in cancer cells is crucial, as it is linked to various cancer subtypes and can provide valuable insights for therapeutic decision-making (Baudrin LG et al. 2018).

To lessen the data sparsity and capture intricate relationships between genes and proteins, we propose a deep learning model, RCoxNet, which uses gene mutation data and provides transparent biological insights for survival analysis, contributing by explicitly incorporating protein interactions and RWR into the model and predict the patient’s survival outcome using clinical features. In this work, we used cBioportal mutation data for four cancer types to obtain gene interaction information using the RWR algorithm. As a result, the RWR score matrix is obtained, which is used as an input layer to the RCoxNet model. The model produces a predictive score called prognostic index (PI) that signifies the patient’s likelihood of survival. The accuracy of our model with the existing models is assessed using the concordance index (C-index). Compared with other current models, our findings illustrate that our suggested framework delivers reliable predictions across four types of cancer.

## 2 Methods

### 2.1 Overview of RCoxNet

RCoxNet is a novel deep learning framework designed for cancer survival prediction, integrating the RWR algorithm with TCGA mutation, clinical and patient data from cBioportal. The framework is structured into four key components: (1) **Gene layer**: This layer uses a matrix of RWR scores, which is calculated from a protein-protein interaction network derived from StringDB along with the RWR algorithm applied using mutated genes as seed nodes. The RWR score layer consists of nodes corresponding to the number of genes identified within the specific cancer context. Each node represents a distinct gene in the input RWR vector, ensuring that each gene’s network-based relevance is captured within the model, (2) **Multiple hidden layers**: The proposed architecture consists of three hidden layers where the number of hidden nodes decreasing progressively from 1^st^ layer (860 nodes) to 3^rd^ layer (30 nodes) passing through an intermediate layer of 100 nodes. These layers capture non-linear relationships within the input data which learn complex patterns and gene interactions. The ’tanh’ activation function is applied within these layers, (3) **Clinical layer**: This layer incorporates clinical covariates such as age, TMB and MSI, allowing the model to integrate key clinical parameters alongside the gene mutation information, and (4) **Cox layer**: This final output layer contains a single node that generates a linear predictor—referred to as the Prognostic Index (PI)—by integrating mutation and clinical data. The PI is fed into a Cox Proportional Hazards model, which accurately stratified patients into high- and low-risk groups based on survival outcomes. The workflow of RCoxNet is illustrated in Fig 1.

**Fig 1.**
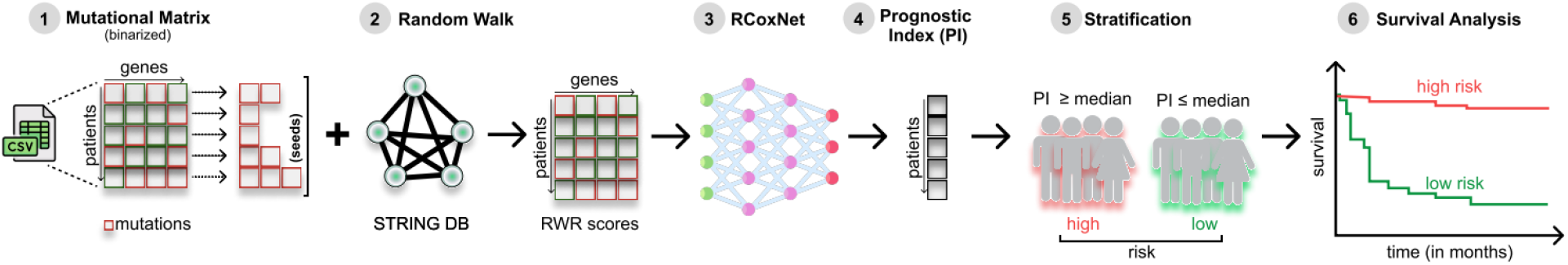
Structure and workflow of proposed model, RCoxNet. Initially, seeds are generated from a mutational matrix (1). Protein-protein interaction list and seeds are combined to serve as an input for the RWR algorithm, resulting in the RWR score matrix (2). The RWR score matrix is then partitioned into the train, validation, and test datasets, integrating them into the RCoxNet model (3). The model yields a prognostic index (PI) for all patients (4). Patients are stratified into high-risk and low-risk based on the median PI value (5). Finally, survival analysis is performed for high-risk and low-risk patients, yielding a corresponding P-value is obtained.

### 2.2 Objective Function

To perform Cox-proportional hazards regression in the Cox layer, RCoxNet defines its objective function by leveraging the average negative log partial likelihood (NLP) alongside L2 regularization to prevent overfitting in the neural network. In survival analysis, particularly when handling censored data, the NLP is a key metric used to estimate model parameters. By minimizing the NLP, the model identifies the coefficient values that best represent the observed survival times and event occurrences. The inclusion of L2 regularization mitigates the risk of overfitting by penalizing excessively large coefficient magnitudes, thus promoting model generalizability.

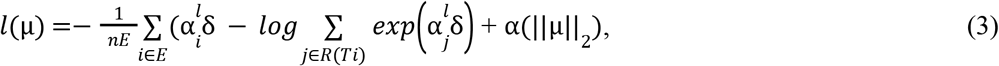

In this equation, µ={δ, *W*} are the set of parameters that needs to be optimized where δ is Cox proportional hazard coefficient that is the weight between last hidden layer and Cox layer for output. *W* is a union of weight matrices across the architecture before the Cox layer. 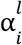 is the very last hidden layer output with the clinical information. *n*_*e*_ is a total number of uncensored events & α is the regularization hyper-parameter. R(T_i_) = { i|T_i_ ≥ *t*} is a set of all samples at risk of failure at time t.

### 2.3 Random Walk with Restart (RWR) algorithm

The Random Walk with Restart (RWR) algorithm is a robust method for investigating gene interactions within gene networks, particularly in functional genomics, disease gene prediction, and biomarker identification. In the RWR method, known genes associated with a specific function or disease, referred to as seeds, serve as the initial nodes of the network. The algorithm evaluates neighboring genes based on their proximity to these seeds, assigning higher scores to genes that are closer to the seeds. We employed an α value of 0.3 as the restart probability, i.e., the random walker has a 30% chance of returning to the seed node at each step. Higher α (e.g., 0.7 to 0.9) is used when the focus is on nodes that are closely related to the seed nodes whereas lower α (e.g., 0.1 to 0.3) encourages more exploration of the network, leading the random walker to visit nodes further from the seed nodes. To integrate RWR into our research, we utilized the RandomWalkRestartMH package in R, which calculates gene scores from protein networks (Valdeolivas A et al., 2019). The application of the RWR algorithm on a monoplex network involves several critical steps, including the computation of the adjacency matrix, its normalization, and the definition of seed nodes. RWR is instrumental in analyzing gene interactions and predicting candidate disease genes by prioritizing genes based on their connectivity to known disease-associated genes within a biological network. The algorithm explores extensive and complex networks of biological molecules, such as protein-protein interactions and co-expression associations. The ability of the RWR algorithm to effectively navigate and prioritize genes within intricate biological networks highlights its significance as a powerful tool in computational biology, offering substantial potential for advancing our understanding of gene function and disease pathogenesis.

i. **Adjacency Matrix (A):** Given a monoplex network represented by a set of nodes V and edges E, the adjacency matrix A is a square matrix where each element A(i, j) is numerical and represents the weight or connection strength between node i and node j. If there is an edge between node i and node j, A(i, j) is a non-zero value reflecting the strength of that connection; otherwise, if there is no edge, (A(i, j) = 0). The higher the value of A(i, j), the stronger the node connection. In our case weight represents protein-protein interaction score.
ii. **Normalization of Adjacency Matrix (P)**: The normalized matrix P is calculated by dividing each connection strength A(i,j) from node i to its neighbors by the sum of the connection strengths from node i to all other nodes. This normalization ensures that the probabilities of transitioning from node i to its neighbors sum up to 1, making it suitable for random walk simulations.

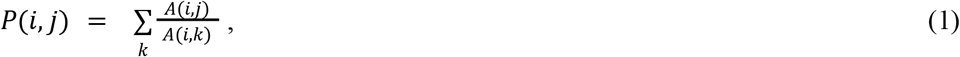

where *P*(*i,j*) is the normalized connection strength from node *i* to node *j, A*(*i,j*) is the original connection strength from node *i* to node *j* in the adjacency matrix, 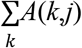 is the sum of connection strengths from node *i* to all other nodes, i.e., the sum of the elements in the column *j* of the adjacency matrix.

#### Random Walk with Restart (RWR) Equation

The RWR algorithm involves iteratively propagating information through the network. The probability distribution of a random walker at each step is updated using the following equation:

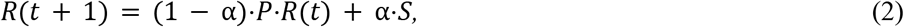

*R*(*t*) is the probability distribution of nodes at time step t. *α* is the restart probability, a parameter between 0 and 1.Choosing the appropriate value for the restart probability α in the Random Walk with Restart (RWR) algorithm is crucial. This is useful when you want to capture more global network information and are interested in relationships that are not just local to the seed nodes. *P* is the normalized adjacency matrix. *S* is a seed vector representing the initial probability distribution of seed nodes.

### 2.4 Data download and preprocessing

For model development, we collected clinical, patient, and TCGA mutation data for Breast Invasive Carcinoma (BRCA, 1084 samples), Lung Adenocarcinoma (LUNG, 566 samples), Glioblastoma Multiforme (GBM, 592 samples), and Ovarian Serous Cystadenocarcinoma (OV, 585 samples) cancers from the cBioPortal database (https://www.cbioportal.org/). These cancers were selected due to their high prevalence and significant mortality rates, representing major clinical challenges in oncology. The datasets were merged based on a common attribute, Patient_id. Following data preprocessing and merging, including the exclusion of missing data, a total of BRCA (1009 samples), LUNG (491 samples), OV(345 Samples), and GBM (264 samples) were retained for further analysis. For each cancer type, we extracted the clinical information for analysis: mutated gene names, OS_Months (Overall Survival Months), OS_Status (Overall Survival Status), Age, MSI_SCORE_MANTIS, and TMB_NONSYNONYMOUS. Notably, the MSI scores for OV were excluded from the analysis due to a substantial amount of missing values. To incorporate network-based features, we downloaded the human protein interaction network from STRING db (Version: V11.5) (https://string-db.org/cgi/download.pl) and converted it into a gene interaction network. The Ensembl Protein IDs were converted to Gene Symbols using the BioDBNet (Version: 2.1) REST API. Additionally, we incorporated driver mutation data from the COSMIC database by downloading the *Cosmic_MutantCensus_Tsv_v99_GRCh37*.*tar* file, which contains 739 genes from the Cancer Gene Census. The COSMIC gene sets for each cancer type comprised 549 genes (BRCA), 512 genes (OV), 519 genes (GBM), and 551 genes (LUNG). These driver genes were mapped onto the gene lists generated through the RWR algorithm to improve model accuracy and gain functional insights. For the RWR computation, we used the mutated genes as seed nodes to initiate a random walk within the STRING protein-protein interaction (PPI) network. RWR scores were calculated for all genes in the network, and for each cancer type, we focused on extracting scores for genes overlapping with the COSMIC gene sets. The resulting RWR scores were integrated into the dataset, where each row represented a patient and each column corresponded to a gene from the COSMIC database. The data integration process involved merging the clinical features (e.g., age, MSI Score, TMB Score) with the RWR gene scores to create a comprehensive input matrix. This matrix, containing both clinical and network-based features, served as the input for our model to predict patient survival outcomes.

### 2.5 Model Training & Hyperparameter Tuning

The dataset was partitioned into 70%, 10%, and 20% for training, validation, and testing, respectively (Table S1). The Adam optimizer was employed for model training, while early stopping was implemented to halt training once the validation loss ceased to improve. Specifications used for testing purposes are RAM: 8+ GB CPU: 2 cores GPU: 15360MiB. A grid search approach was utilized for hyperparameter optimization, exploring exhaustive combinations of L2 regularization (0.1, 0.01, 0.005, 0.001) and learning rates (0.03, 0.01, 0.001, 0.0075) to identify the optimal set. During this process, the model was trained and evaluated for each parameter combination, and the set yielding the lowest validation loss was selected. The final model was then built with these optimized hyperparameters to ensure reproducibility of model performance.

### 2.6 Survival Analysis

Kaplan–Meier (KM) curves were employed to visualize the survival outcomes of the predicted high- and low-risk groups. Survival analysis was conducted using the Python package lifelines (Version 0.28.0). Specifically, the KaplanMeierFitter function from the lifelines library was utilized to fit Kaplan–Meier survival curves for both risk groups, and the resulting survival distributions were plotted to illustrate differences in patient outcomes.

## 3 Results

### 3.1 RCoxNet model on mutation, clinical and patient data

The structure of RCoxNet was presented in Fig 1 and the “Methods” section. In general, the RCoxNet model conducts survival prediction in two steps: (1) RWR algorithm with protein-protein interaction score extracts higher-order relational features from mutation, clinical and patient data; (2) Combination of all molecular and clinical features as input and computes prognostic index (PI) for each patient. The correlation between random walk scores and survival rates allows for the identification of prognostic genes linked to survival outcomes. The utilization of random walk scores for stratifying patients results in more homogeneous groupings, revealing unique survival patterns. Following the execution of our model, the PI is computed and applied to categorize patients into two groups - high-risk and low-risk groups. Patients with a PI greater than the median value (PI > median value) are categorized as high-risk, indicating a worse prognosis and a higher likelihood of the event (e.g., death) occurring within a given timeframe. Conversely, patients with a PI less than the median value (PI < median value) are classified as low-risk, signifying a more favorable prognosis and a reduced risk of the event.

Fig 2 illustrates the Kaplan-Meier plot, demonstrating p-values below 0.05 for all cancer types, providing strong evidence against the null hypothesis. For GBM, as the median PI value is 2.824, patients with a PI exceeding this threshold are categorized as high-risk, indicating a worse prognosis, while those with a PI below 2.824 are classified as low-risk, reflecting a better chance of survival. Similarly, In the case of OV, a median PI of 1.68 stratifies patients into high-risk (PI > 1.68), associated with a worse prognosis, and low-risk (PI < 1.68), indicating improved survival prospects. For LUNG, a PI threshold of 0.529 distinguishes high-risk patients (PI > 0.529), with a higher likelihood of adverse outcomes, from low-risk patients (PI < 0.529), who exhibit a reduced mortality risk. Finally, for BRCA, a median PI of 1.99 is used to classify patients, with those exceeding this value considered high-risk, indicating a poorer prognosis, while those below this threshold are classified as low-risk, suggesting better survival outcomes.

**Fig 2.**
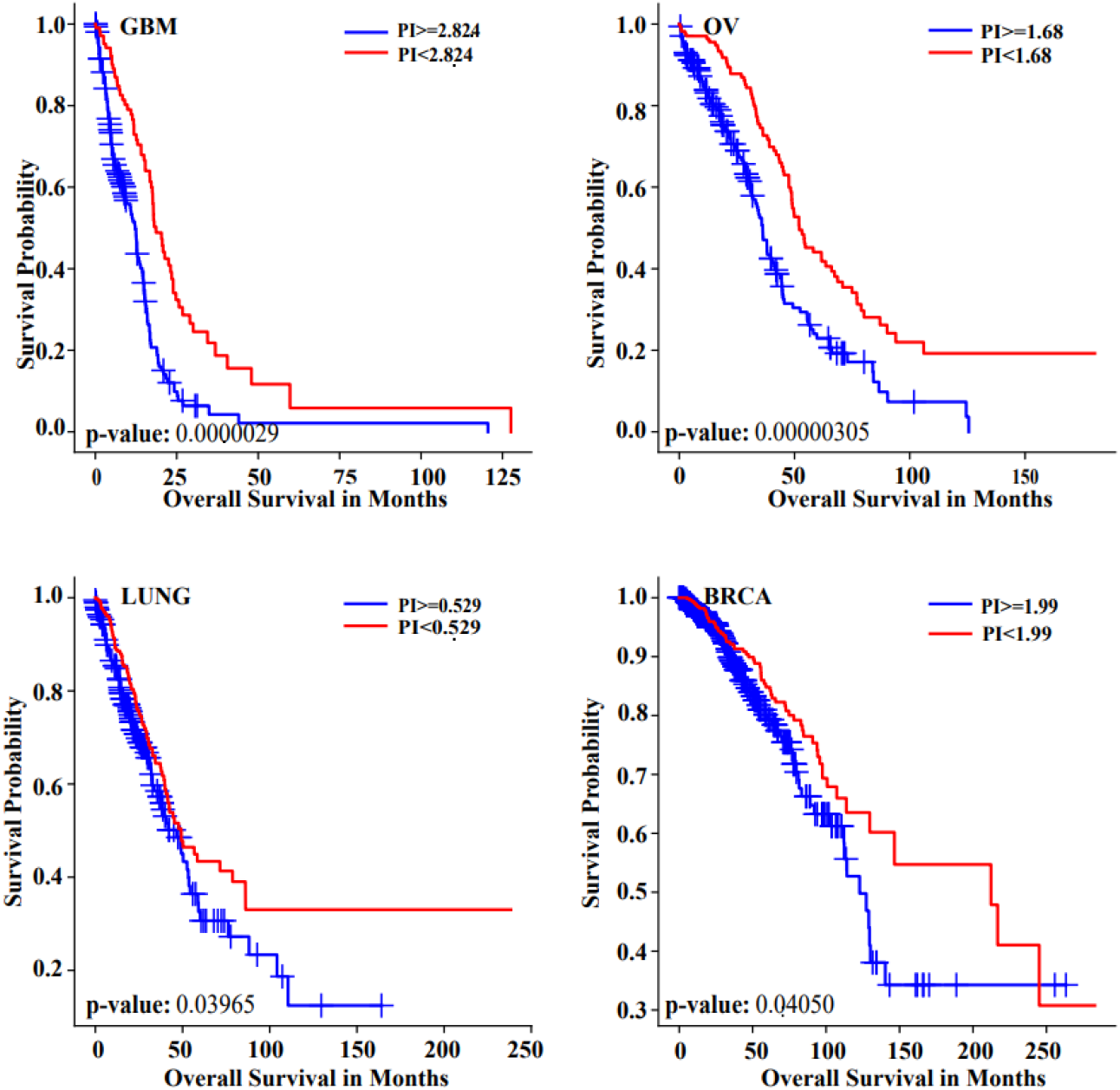
Kaplan Meier curves delineating the Prognostic Index across diverse cancer types reveal distinct trajectories. These graphical representations elucidate a compelling association: elevated PI correlates with diminished survival, manifesting as poor prognostic outcomes, while lower PI values correspond with heightened survival, indicative of a more favorable prognostic profile.

### 3.2 Combining clinically relevant molecular features enhances survival prediction performance

Apart from the features such as scores obtained from the RWR score matrix, clinical data (e.g., age, TMB, MSI) has been combined to enhance survival prediction performance. To confirm this hypothesis on this combined data, a univariate Cox model was employed to validate its statistical significance (Fig 3). Patients underwent stratification based on TMB score, MSI score, and age, unveiling the robust predictive capacity of these covariates. This categorization facilitated a distinct segregation between high-risk and low-risk cohorts, showcasing the efficacy of this classification paradigm. The complex relationship between tumor mutational burden (TMB) and survival rate varies depending on the cancer type and other clinical factors (Wang P et al., 2021). MSI is distinctly associated with certain cancers. Figure 3a portrays the survival curve in breast cancer revealing that high MSI corresponds to diminished survival, while low MSI correlates with heightened survival (Vidula N et al., 2022). As delineated in Figure 3b, breast cancer exemplifies improved survival rates with elevated TMB due to immune system recognition of cancer cells, consequently resulting in diminished survival rates when TMB is low (Ke L et al., 2022). Although TMB and MSI are distinct markers, occasional overlap can occur. Some tumors classified as MSI-high may also exhibit elevated TMB. It is imperative to underscore that these markers are not interchangeable, and a tumor can manifest as either TMB-high and MSI-stable or vice versa. The supplementary figures, denoted as Figure S1, S2 and S3, depict the plots corresponding to age, TMB, and MSI for various cancer types. The association between advancing age and heightened likelihood of mortality persists as a salient observation. The continuous variable of patients’ age correlates independently with overall survival even after meticulous adjustment for covariates. Figure 3c elucidates this correlation through a survival curve, illustrating lower survival rates among older individuals in contrast to higher survival rates observed in younger age groups, contingent upon the prognostic index (PI) (Jackson EB et al., 2023). Although many studies indicate that high TMB is associated with better survival in cancer patients, this conclusion is not applicable to all cancer types.

**Fig 3.**
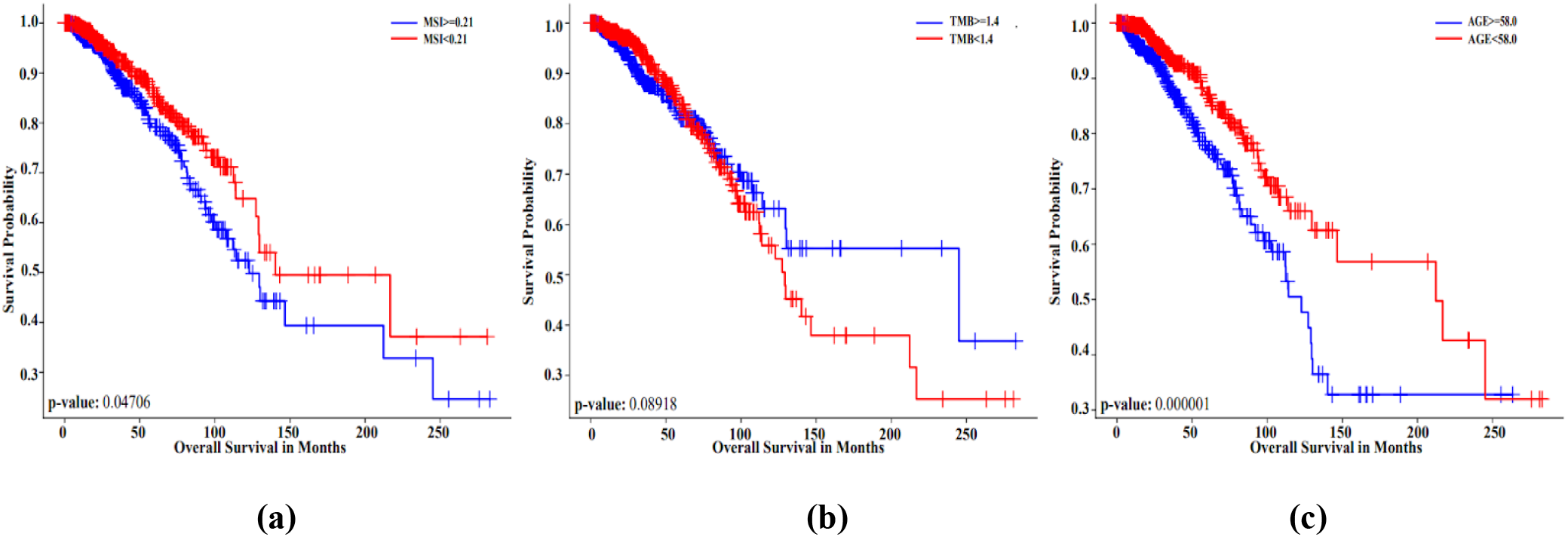
Survival curves of all clinical features for BRCA cancer, (a) MSI, (b) TMB, (c) Age

### 3.3 RCoxNet model shows superior performance as compared to state-of-the-art deep learning approaches

We applied and adapted other current state-of-the-art deep learning methods, DeepSurv (Katzman JL et al. 2018) and DeepHit (Lee C et al. 2018) with the same hyperparameters as mentioned earlier for RCoxNet and compared their performance. For measuring predictive performance with censored data, C-index, a rank-correlation method that counts concordant pairs between the predicted score and observed survival time, assigns a score between 0 and 1, where 1 indicates an ideal prediction, and 0.5 indicates a random prediction. DeepHit is a neural network model trained on the Probability Mass Function (PMF) of a discrete Cox model whereas DeepSurv is a deep feed-forward neural network that forecasts how a patient’s covariates influence their hazard rate. For implementation of DeepSurv and DeepHit, pycox (GitHub - havakv/pycox: Survival analysis with PyTorch) package has been utilized. Pycox (Version: 0.2.3) is a python package for survival analysis and time-to-event prediction with pytorch (Version: 2.0.1, CUDA Version: 12.1), built on the torchtuples package for training PyTorch models.

Our model underwent rigorous validation against prominent deep neural network models, DeepSurv and DeepHit, both intricately tailored for the domain of cancer survival prediction. RCoxNet demonstrated a notably elevated average concordance index (C-index) of 0.62 ± 0.05 across four distinct cancer types. In contrast, DeepHit claimed the second position with an average C-index of 0.54 ± 0.0332, while DeepSurv exhibited a comparatively diminished average C-index of 0.50 ± 0.12 within the triad of models considered. The outcomes of experimental analyses unequivocally substantiate the heightened efficacy and superior performance of RCoxNet when juxtaposed with contemporary state-of-the-art deep neural network models (Table 1, Fig 4).

**Table 1.** C-index of existing deep neural network models along with RCoxNet across four cancer types.

|  | **RCoxNet with existing deep neural network models** |  |  |  |
| --- | --- | --- | --- | --- |
|  |  | **DeepSurv** | **DeepHit** | **RCoxNet** |
| **Cancer types** | **OV** | 0.42 | 0.49 | **0.60** |
|  | **BRCA** | 0.50 | 0.49 | **0.66** |
|  | **LUNG** | 0.46 | 0.51 | **0.57** |
|  | **GBM** | 0.39 | 0.54 | **0.65** |

**Fig 4.**
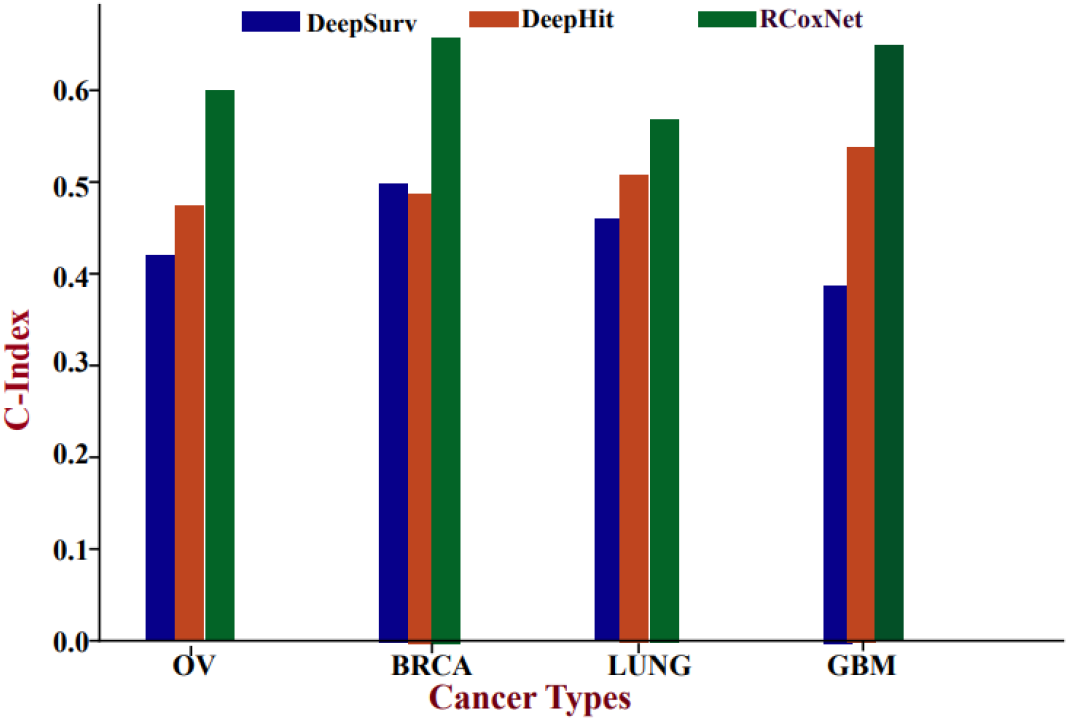
Comparison of concordance index (C-index) of RCoxNet model with existing deep learning models, DeepSurv and DeepHit, across four types of cancer.

## 4 Discussion

In this paper, RCoxNet, an innovative deep learning architecture designed for the analysis of patient survival outcomes utilizing TCGA mutation data in conjunction with the RWR algorithm. While previous efforts have tried to use mutation data with deep learning models for accurate predictions, these models still haven’t provided a complete biological interpretation. Our proposed model RCoxNet confers distinct advantages over prevailing models by leveraging mutation data in tandem with the RWR algorithm. The incorporation of the RWR layer, alongside clinical layers encompassing age, TMB and MSI, within the RCoxNet model significantly contributes to survival analysis, delineating its efficacy as a prognostic indicator. This approach, culminating in the computation of PI, facilitates a nuanced understanding of patient prognosis and survival rates. The inclusion of age, TMB and MSI as covariates in the model imparts a substantial impact on survival rates. In summary, age, TMB, and MSI emerge as robust predictors of survival, establishing themselves as valuable prognostic biomarkers in the realm of cancer. Thus, this methodology holds promise as a pivotal tool for personalized treatment strategies, making noteworthy contributions to patient survival rates beyond the confines of conventional survival-based models.

Despite our findings, our study still has some limitations. First, the number of cases included in this study is limited, and more data are needed to confirm our conclusions. Second, the standard of high TMB in some studies is not clear; thus, we did not discuss the specific critical value of TMB as a predictive marker. Finally, each case might have received different treatments before being included in this study, which may lead to different therapeutic effects.

## Data and materials availability

The code and data can be obtained from https://github.com/stutimishra7/Survival-Analysis.

## Authors contributions

Conceptualization: DS. Methodology: DS, SK, SG. Investigation: SK, SG, SP. Visualization: SK, SG, SP. Supervision: DS. Writing – original draft: DS, SK, SG, SP. Writing – review & editing: DS, SK, SG, SP, PG, GA.

## References

Baudrin LG, Deleuze JF, How-Kit A. Molecular and computational methods for the detection of microsatellite instability in cancer. Frontiers in oncology. 2018 Dec 12;8:621.

Chakrabarti S, Bucheit L, Starr JS, Innis-Shelton R, Shergill A, Dada H, Resta R, Wagner S, Fei N, Kasi PM. Detection of microsatellite instability-high (MSI-H) by liquid biopsy predicts robust and durable response to immunotherapy in patients with pancreatic cancer. Journal for immunotherapy of cancer. 2022 Jun 1;10(6):e004485.

Ching T, Zhu X, Garmire LX. Cox-nnet: an artificial neural network method for prognosis prediction of high-throughput omics data. PLoS computational biology. 2018 Apr 10;14(4):e1006076.

Cui S, Feng J, Tang X, Lou S, Guo W, Xiao X, Li S, Chen X, Huan Y, Zhou Y, Xiao L. The prognostic value of tumor mutation burden (TMB) and its relationship with immune infiltration in breast cancer patients. European Journal of Medical Research. 2023 Dec;28(1):1–1.

Dwivedi B, Mumme H, Satpathy S, Bhasin SS, Bhasin M. Survival Genie, a web platform for survival analysis across pediatric and adult cancers. Scientific Reports. 2022 Feb 23;12(1):3069.

Ghosh Roy G, Geard N, Verspoor K, He S. MPVNN: Mutated Pathway Visible Neural Network architecture for interpretable prediction of cancer-specific survival risk. Bioinformatics. 2022 Nov 15;38(22):5026–32.

Huang Z, Johnson TS, Han Z, Helm B, Cao S, Zhang C, Salama P, Rizkalla M, Yu CY, Cheng J, Xiang S. Deep learning-based cancer survival prognosis from RNA-seq data: approaches and evaluations. BMC medical genomics. 2020 Apr;13:1–2.

Huan T, Wu X, Bai Z, Chen JY. Seed-weighted random walk ranking for cancer biomarker prioritisation: a case study in leukaemia. International Journal of Data Mining and Bioinformatics. 2014 Jan 1;9(2):135–48.

Jackson EB, Gondara L, Speers C, Diocee R, Nichol AM, Lohrisch C, Gelmon KA. Does age affect outcome with breast cancer?. The Breast. 2023 Jun 2.

Katzman JL, Shaham U, Cloninger A, Bates J, Jiang T, Kluger Y. DeepSurv: personalized treatment recommender system using a Cox proportional hazards deep neural network. BMC medical research methodology. 2018 Dec;18(1):1–2.

Ke L, Li S, Cui H. The prognostic role of tumor mutation burden on survival of breast cancer: a systematic review and meta-analysis. BMC cancer. 2022 Dec;22(1):1–2.

Lee C, Zame W, Yoon J, Van Der Schaar M. Deephit: A deep learning approach to survival analysis with competing risks. InProceedings of the AAAI conference on artificial intelligence 2018 Apr 26 (Vol. 32, No. 1).

Liu J, Islam MT, Sang S, Qiu L, Xing L. Biology-aware mutation-based deep learning for outcome prediction of cancer immunotherapy with immune checkpoint inhibitors. NPJ Precision Oncology. 2023 Nov 6;7(1):117.

Liu WX, Shi M, Su H, Wang Y, He X, Xu LM, Yuan ZY, Zhang LL, Wu G, Qu BL, Qian LT. Effect of age as a continuous variable on survival outcomes and treatment selection in patients with extranodal nasal-type NK/T-cell lymphoma from the China Lymphoma Collaborative Group (CLCG). Aging (albany NY). 2019 Oct 10;11(19):8463.

Ma X, Zhang Y, Wang S, Wei H, Yu J. Immune checkpoint inhibitor (ICI) combination therapy compared to monotherapy in advanced solid cancer: A systematic review. Journal of Cancer. 2021;12(5):1318.

Makrooni MA, O’Sullivan B, Seoighe C. Bias and inconsistency in the estimation of tumour mutation burden. BMC cancer. 2022 Dec;22(1):1–9.

Morel LO, Derangère V, Arnould L, Ladoire S, Vinçon N. Preliminary evaluation of deep learning for first-line diagnostic prediction of tumor mutational status. Scientific Reports. 2023 Apr 28;13(1):6927.

Piña-Sánchez P, Chavez-Gonzalez A, Ruiz-Tachiquin M, Vadillo E, Monroy-Garcia A, Montesinos JJ, Grajales R, Gutierrez de la Barrera M, Mayani H. Cancer biology, epidemiology, and treatment in the 21st century: Current status and future challenges from a biomedical perspective. Cancer Control. 2021 Sep 13;28:10732748211038735.

Ramirez R, Chiu YC, Zhang S, Ramirez J, Chen Y, Huang Y, Jin YF. Prediction and interpretation of cancer survival using graph convolution neural networks. Methods. 2021 Aug 1;192:120–30.

Schober P, Vetter TR. Survival analysis and interpretation of time-to-event data: the tortoise and the hare. Anesthesia and analgesia. 2018 Sep;127(3):792.

Sha D, Jin Z, Budczies J, Kluck K, Stenzinger A, Sinicrope FA. Tumor mutational burden as a predictive biomarker in solid tumors. Cancer discovery. 2020 Dec 1;10(12):1808–25.

Sherman MA, Yaari AU, Priebe O, Dietlein F, Loh PR, Berger B. Genome-wide mapping of somatic mutation rates uncovers drivers of cancer. Nature Biotechnology. 2022 Nov;40(11):1634–43.

Strickler JH, Hanks BA, Khasraw M. Tumor mutational burden as a predictor of immunotherapy response: is more always better?. Clinical Cancer Research. 2021 Mar 1;27(5):1236–41.

Sun B, Chen L. Interpretable deep learning for improving cancer patient survival based on personal transcriptomes. Scientific Reports. 2023 Jul 13;13(1):11344.

Valdeolivas A, Tichit L, Navarro C, Perrin S, Odelin G, Levy N, Cau P, Remy E, Baudot A. Random walk with restart on multiplex and heterogeneous biological networks. Bioinformatics. 2019 Feb 1;35(3):497–505.

Vidula N, Lipman A, Kato S, Weipert C, Hesler K, Azzi G, Elkhanany A, Juric D, Rodriguez E, Faulkner C, Makhlouf P. Detection of microsatellite instability high (MSI-H) status by targeted plasma-based genotyping in metastatic breast cancer. NPJ Breast Cancer. 2022 Nov 4;8(1):117.

Wang J, Chen N, Guo J, Xu X, Liu L, Yi Z. SurvNet: a novel deep neural network for lung cancer survival analysis with missing values. Frontiers in Oncology. 2021 Jan 20;10:588990.

Wang P, Chen Y, Wang C. Beyond tumor mutation burden: tumor neoantigen burden as a biomarker for immunotherapy and other types of therapy. Frontiers in oncology. 2021 Apr 29;11:672677.

Yousefi S, Amrollahi F, Amgad M, Dong C, Lewis JE, Song C, Gutman DA, Halani SH, Velazquez Vega JE, Brat DJ, Cooper LA. Predicting clinical outcomes from large scale cancer genomic profiles with deep survival models. Scientific reports. 2017 Sep 15;7(1):11707.

Zheng W, Pu M, Li X, Du Z, Jin S, Li X, Zhou J, Zhang Y. Deep learning model accurately classifies metastatic tumors from primary tumors based on mutational signatures. Scientific Reports. 2023 May 30;13(1):8752.

