## Supplementary Material for "RCoxNet: deep learning framework for enhanced cancer survival prediction integrating random walk with restart with mutation and clinical data"

**Supplementary materials: RCoxNet: Network-diffusion powered Deep Cox Proportional Hazard model for improved survival prediction based on cancer mutation data**

Stuti Kumari^2^*^#^*, Sakshi Gujral^2^*^#^*, Smruti Panda^1^, Prashant Gupta^4^, Gaurav Ahuja^1,3^, Debarka Sengupta^1,2,3+^

1. Department of Computational Biology, Indraprastha Institute of Information Technology-Delhi (IIIT-Delhi), Okhla, Phase III, New Delhi-110020, India
2. Department of Computer Science and Engineering, Indraprastha Institute of Information Technology-Delhi (IIIT-Delhi), Okhla, Phase III, New Delhi-110020, India
3. Center for Artificial Intelligence, Indraprastha Institute of Information Technology-Delhi (IIIT-Delhi), Okhla, Phase III, New Delhi-110020, India
4. Wellcome Sanger Institute, Wellcome Trust Genome Campus, Hinxton, Saffron Walden CB10 1RQ, UK

*^#^Equal contribution*

| **Cancer Types** | **Train** | **Test** | **Validation** | **Total** |
| --- | --- | --- | --- | --- |
| BRCA | 706 | 203 | 102 | 1009 |
| LUNG | 343 | 98 | 50 | 491 |
| GBM | 184 | 53 | 27 | 264 |
| OV | 242 | 69 | 35 | 346 |
| Total | 1475 | 423 | 214 | 2110 |

**Table S1.** Distribution of samples across four cancer types & splits

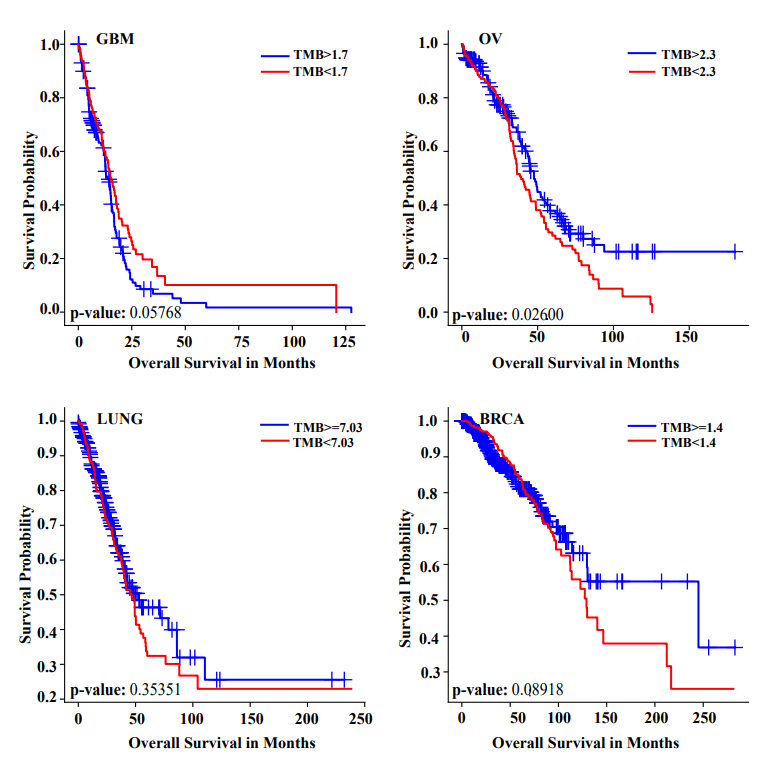

**Fig S1.** Survival curves of Tumor Mutational Burden (TMB) for all cancer types

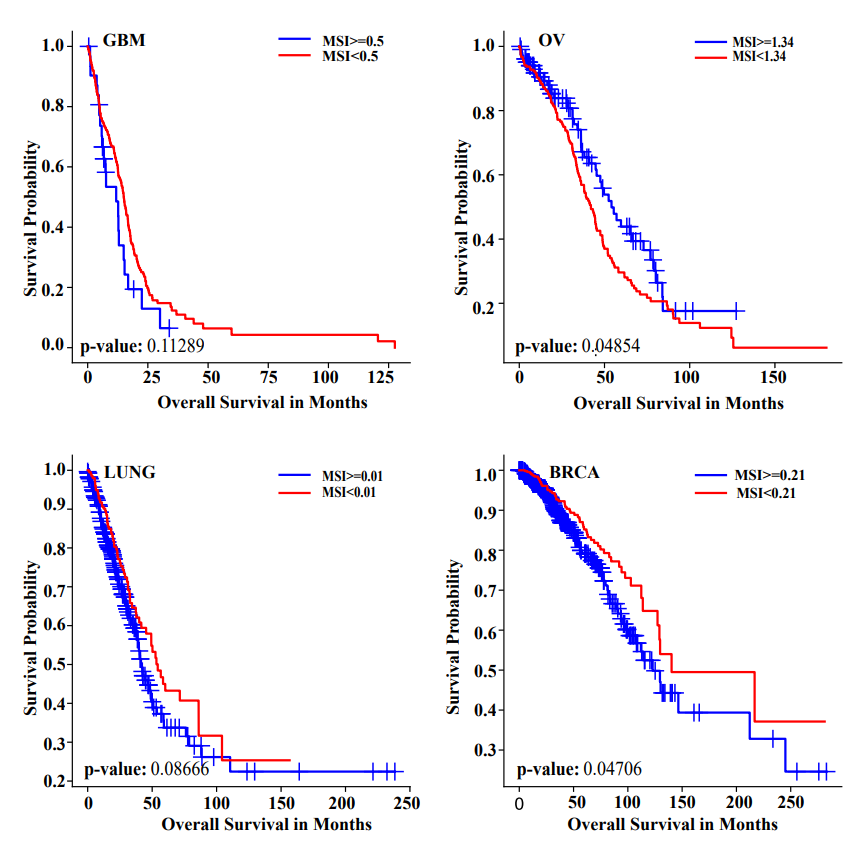

**Fig S2.** Survival curves of Microsatellite Instability (MSI) for all cancer types

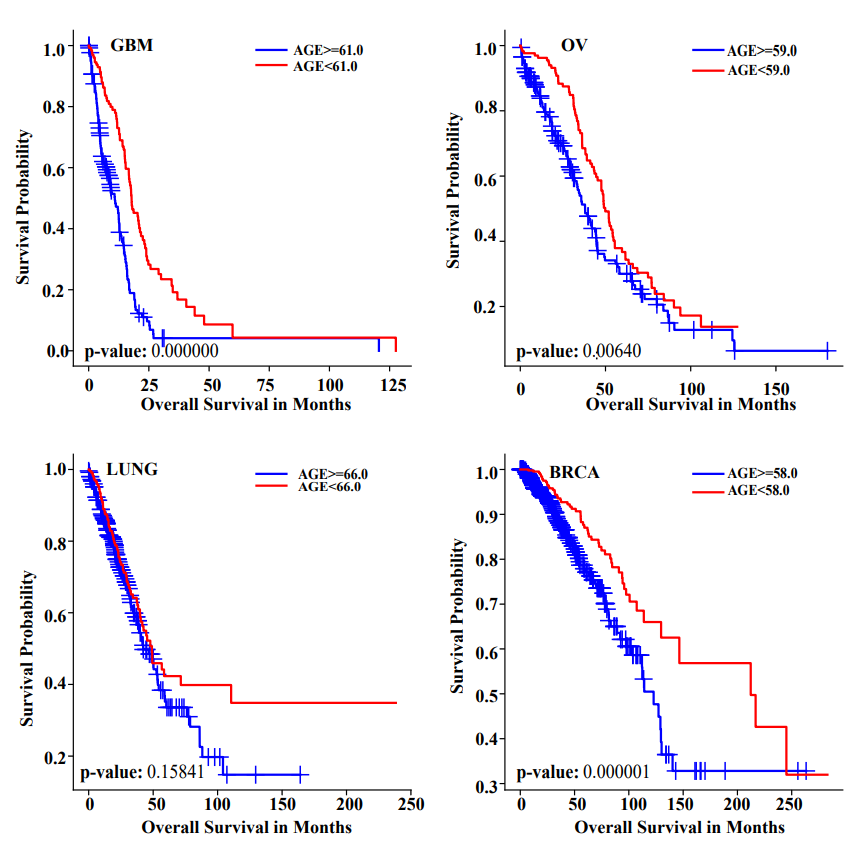

**Fig S3.** Survival curves of age for all cancer types

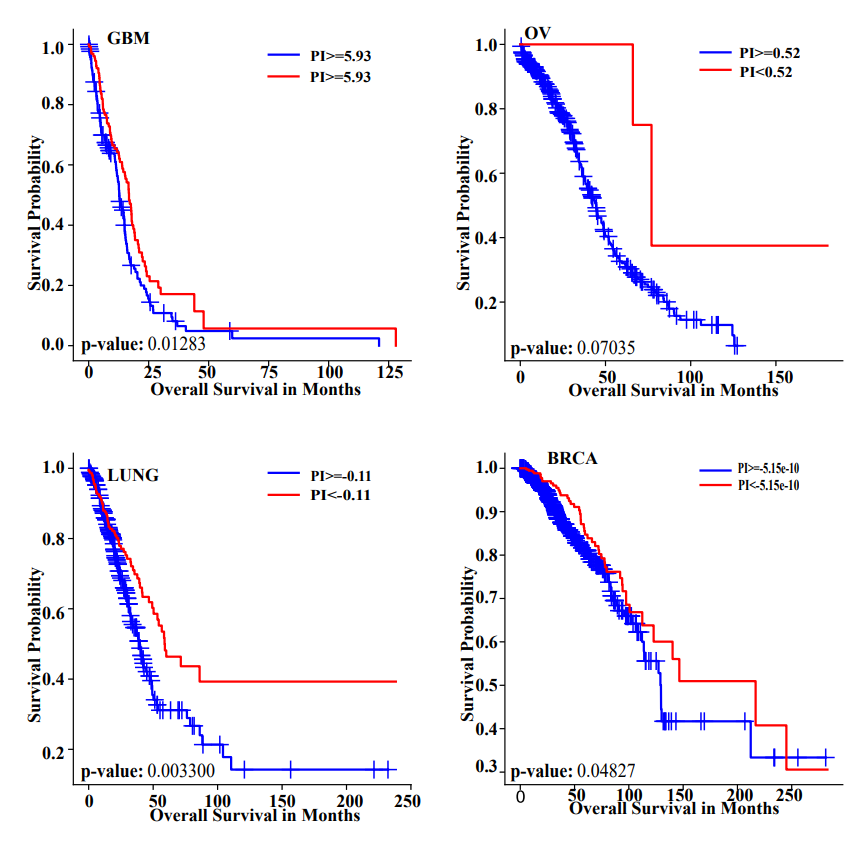

**Fig S4.** Survival curves of RWR for all cancer types
